# Electrophysiological Effects Of Kappa-Opioid Analgesic, RU-1205, Using Machine Learning Methods

**DOI:** 10.1101/2024.02.11.579812

**Authors:** Konstantin Y. Kalitin, Olga Y. Mukha, Alexander A. Spasov

## Abstract

This study focuses on RU-1205, a new kappa-opioid agonist exhibiting analgesic effect without causing dysphoric or aversive reactions. It is assumed that the absence of dysphoric or aversive effects can be attributed to functional selectivity or it might be due to an additional mechanism of action that involves blocking the p38 mitogen-activated protein kinase (MAPK).

**The aim:** of this study was the experimental identification of the mechanisms of action of RU-1205 associated with inhibition of MAPK p38 and functional selectivity at kappa-opioid receptors.

**Materials and methods:** Rats weighing 260-280 g were implanted with chronic cortical and deep electrodes. LFP activity was recorded after intracerebroventricular administration of well-studied reference substances: the selective kappa-opioid agonist U-50488 at a dose of 100 μg; the MAPK p38 blocker SB203580 at a dose of 1 μg; and the investigational compound RU-1205 at 350 μg. The weighted phase lag index (WPLI) was calculated. Subsequently, machine learning techniques were employed to reduce dimensionality and extract connectivity features using the principal component analysis method. Finally, signal classification was conducted using models based on Gaussian processes.

By applying the patch-clamp technique in the whole-cell configuration, the spike activity of pyramidal neurons in the basolateral amygdala was studied. The neurons were identified by their accommodation properties. After local perfusion of the test compounds, 3 dose-response curves were obtained for: (1) U-50488 at concentrations ranging from 0.001 to 10 μM; (2) combinations of U-50488 (0.001–10 μM) and RU-1205 (10 μM); and (3) combinations of U-50488 (0.01–10 μM) and RU-1205 (100 μM).

**Results:** The developed models were able to classify the compound RU-1205 as a «non-inhibitor» of MAPK p38 with a probability of 0.89. The results obtained were confirmed in patch clamp experiments on acute brain slices, where it was demonstrated that U-50488 statistically significantly increases the spike activity of pyramidal neurons in the basolateral amygdala (p <0.05) and RU-1205 interacts with U-50488, suppressing its effect on the spike activity of neurons.

**Conclusions:** The findings suggest that compound RU-1205 displays properties consistent with a functional kappa opioid receptor agonist and does not have a significant effect on MAPK p38. The study demonstrates the possibility of integrating electrophysiological measurements and advanced data analysis methods for a deep understanding of neuronal mechanisms of drug action and underscores the potential for further research in this area.

## INTRODUCTION

Pain is one of the most common and challenging medical issues facing our society [1]. Regardless of its location or nature, whether acute or chronic, the use of analgesics remains a primary method for pain management [2].

Most of the narcotic analgesics used in clinical practice act on the mu-opioid receptor type; however, the use of these drugs is associated with serious side effects, such as the development of dependence, nausea, respiratory depression, and constipation[3]. It was found that agonists of kappa-opioid receptors have an analgesic effect without causing adverse reactions in the respiratory and gastrointestinal tract, and also have a low abuse potential [4]. However, their application is constrained due to aversive effects such as depression, dysphoria, and hallucinations [5].

Kappa-opioid receptors are abundant in the central and peripheral nervous systems. They are involved in the regulation of pain, mood, behavior, motor activity, and perform neuroprotective and neuroendocrine functions [5, 6]. Active conformations of the receptor are required for the recruitment of effector molecules, the main of which are G-proteins and beta-arrestins. It is assumed that the analgesic effect of kappa-opioid agonists is associated with activation of G-proteins and subsequent inhibition of adenylate cyclase, increasing potassium and decreasing calcium conductivity [7]. Activation of the beta-arrestin pathway leads to suppression of G-protein signaling, desensitization and internalization of the kappa-opioid receptor, which is one of the reasons for the development of tolerance. In addition, beta-arrestins can initiate p38 protein kinase, which is an important component of the mechanism for the development of aversive side effects [8].

An urgent task is to search for safer kappa-opioid receptor agonists with analgesic activity [9]. Among the solutions to this problem, the following areas can be identified: 1) the creation of drugs that have only a peripheral effect, that is, they do not penetrate the blood-brain barrier (BBB) and do not act on the central nervous system (CNS); 2) the search for functionally selective substances that activate a specific signaling pathways, such as G-proteins, without involving beta-arrestins [10]. It is assumed that the conformational changes of the receptor upon binding to the agonist determine functional selectivity.

Our research focuses on the new kappa-opioid agonist RU-1205. Unlike classical representatives of this class, the compound RU-1205 exhibits pronounced analgesic, anticonvulsant [11, 12] and neuroprotective [13] effects while not causing dysphoric or aversive effects. This property may be due to functional selectivity, or the presence of an additional mechanism of action that is associated with blocking p38 mitogen-activated protein kinase (MAPK), since it was previously shown that the p38 inhibitor SB203580 can completely eliminate the aversive effect of kappa-opioid agonists [14].

Since aversive effects can manifest at both the cellular and systemic levels of neuronal organization, we employed the patch clamp method and recorded local field potentials (LFP), followed by the calculation of the weighted phase lag index (WPLI) and the application of machine learning (ML) as a tool for identifying the differences and similarities among ligands with well-studied pharmacological properties, such as U-50488 and the p38 MAPK inhibitor SB203580.

### THE AIM

An investigation of the mechanisms of action of RU-1205 associated with inhibition of MAPK p38 and functional selectivity. The results of this study will expand our understanding of the molecular mechanisms underlying the analgesic and side effects of kappa opioid agonists.

## MATERIALS AND METHODS

### Study design

The study was conducted in two main stages. In the first stage, the local field potential (LFP) was recorded after the administration of the test substances, then the weighted phase delay index was calculated, and the resulting data were analyzed using machine learning methods.

During the second stage, the interaction of the compound RU-1205 with U-50488 at the cellular level was studied using the patch clamp method in order to clarify the mechanism of action of RU-1205 and further validate the results of the LFP classification.

### Test compounds

The compounds used in this study were as follows: 9-(2-morpholinoethyl)-2-(4-fluorophenyl) imidazo[1,2-a]benzimidazole – RU-1205 (synthesized at the Research Institute of Physical and Organic Chemistry of the Southern Federal University, Russia); compound U-50488 (Sigma Aldrich, USA); compound SB203580 (Sigma Aldrich, USA).

### Experimental timeline and setting

The study was conducted between May 2023 and September 2023. All experimental procedures were performed in the electrophysiological research laboratory of the Scientific Center for Innovative Drugs of the Volgograd State Medical University of the Ministry of Health of the Russian Federation.

### Compliance with ethical standards

Animal experiments were carried out in accordance with the European Convention for the Protection of Vertebrate Animals Used for Experimental and Other Scientific Purposes, the principles of Good Laboratory Practice (GLP) (GOST 33044-2014, 2021), as well as the ARRIVE (Animal Research: Reporting of In Vivo Experiments). The study was approved by the Local Ethics Committee of the Volgograd State Medical University of the Ministry of Health of Russia (Registration number IRB00005839 IORG0004900, Minutes No. 2022/096 of 21.01.2022).

### Animals

Sixty-two white male rats, weighing 260–280 g, were used in these studies. Animals were subjected to a 12-hour light/dark cycle, the ambient temperature was 22±2°C, with food and water available *ad libitum*. The rats were bred in the vivarium of the Scientific Center for Innovative Drugs at Volgograd State Medical University.

### Surgery

Implantation of stainless steel electrodes (∅ 0.1 mm) was carried out under isoflurane anesthesia (Laboratories Karizoo, S.A., Spain) using an available rodent inhalant anesthesia system (Ugo Basile Veterinary anesthesia workstation 21100, Italy). The electrodes were stereotaxically inserted through burr holes to target sites in accordance with the coordinates relative to bregma.

Cortical electrodes: F – anteroposterior axis (AP)=0.00, mediolateral axis (ML)=2.00; P – AP=-4.08, ML=2.00; O – AP=-7.08, ML=2.00.

Deep electrodes: prelimbic cortex (PrL) – AP=+2.7 mm, ML=0.8 mm, dorsoventral (DV)=3.8 mm; basolateral amygdala (BLA) – AP=-2.8 mm, ML=5–5.3 mm, DV=8.8 mm; hippocampus (Hipp) – AP=-4.9 mm, ML=4.8 mm, DV=6.0 mm; ventral tegmental area (VTA) – AP=-5.2 mm, ML=1.0 mm, DV=8.6 mm; nucleus accumbens (NAc) – AP=+1.8 mm, ML=1.6 mm, DV=7.3 mm.

For the intracerebroventricular (i.c.v.) administration of drugs, a 21-gauge stainless steel guide cannula was implanted into the right lateral ventricle. The cannula was positioned based on the stereotaxic coordinates relative to the bregma: AP=-0.6 mm, ML=1.6 mm, DV=4.0 mm. The intracerebroventricular route of administration was chosen to enhance the bioavailability of substances and to prevent the effects outside the CNS, which could potentially mediate secondary impacts on brain functions and distort the experimental results. Two stainless steel screws and dental polymer (Protacryl-M; Stoma, Ukraine) were used to secure the cannulas and electrodes to the skull surface. After the surgery, animals were individually housed, and antibiotic prophylaxis with ciprofloxacin at a dose of 50 mg/kg intraperitoneally was administered for two days. The postoperative period lasted 5–7 days before the start of signal recording.

### LFP signal recording and administration of studied substances

After adaptation, animals (*n*=30) received one of the following treatments: ACSF 5 μl (control sample), 350 μg of compound RU-1205, 100 μg of compound U-50488, 1 μg of compound SB203580, and a combination of SB203580 and U-50488 (1 μg and 100 μg, respectively) intracerebroventricularly. Intervals of 7 days were maintained between administrations of substances to achieve near-complete elimination of the previous dose (5 fold half-life). Doses for compounds U-50488 [15] and SB203580 [16] were selected based on literature data. Compound RU-1205 was administered at a dose equivalent to the ED80 in pain models. 30 minutes after administration of the compounds, LFP recordings were obtained for 10 minutes. LFP was recorded in a monopolar montage with a common average reference using a laboratory electroencephalograph NVX-36 (MKS, Russia). Signals were digitized at a sampling rate of 500 Hz.

### WPLI analysis

To assess functional connectivity between electrode pairs, a weighted phase delay index was used (Fig. 1). Calculation of WPLI between electrodes was carried out using the MNE Python software package v.1.6.1^1^ (BSD-3-Clause license) for the following frequency ranges: delta 0.5–4 Hz, theta 4–8 Hz, alpha 8–12 Hz, beta 12–30 Hz and gamma 30–50 Hz. The obtained data were analyzed by the principal component analysis method using the Graphpad Prism 10.1 (Dotmatics, USA) with an academic license.

**Figure 1.**
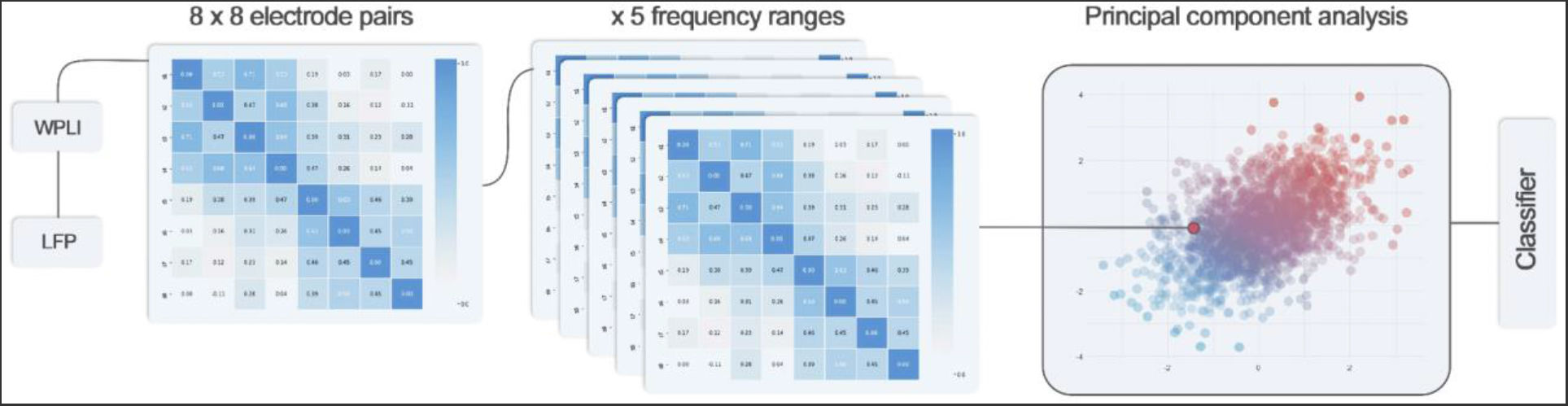
Main steps of data analysis and mathematical modeling. Note: After calculating the weighted phase latency index (WPLI), which reflects connectivity between brain regions, including the cortex, hippocampus, amygdala, ventral tegmental area, prelimbic cortex, and nucleus accumbens (8 × 8 electrodes × 5 rhythms), the data were processed by principal component analysis and then fed as inputs to the classifier (“Gaussian Process Classifier”, scikit-learn).

### Construction of the ML classifier model

To build models based on machine learning algorithms, we used the open-source Python library scikit-learn 1.3.2^2^ (BSD 3-Clause License). Two classification models were established based on Gaussian processes: GPC-BO-v.02.10-5.2308 (GaussianProcessClassifier with parameters 10.0 * RBF(5.0), optimizer=None) and GPC-BO-v.02.10-10.2308 (GaussianProcessClassifier with parameters 10.0 * RBF(10.0), optimizer=None). After training the models, confusion matrices were built and 5-fold cross-validation was performed.

### Patch clamp in the “whole cell” configuration

Coronal brain slices, 500 μm thick and containing the BLA, were cut 2.5–3.5 mm caudal to the bregma using a vibratome (Campden 7000smz-2, UK). Each slice was transferred to a recording chamber, submerged in artificial cerebrospinal fluid (ACSF) maintained at 31±1°C with a superfusion rate of 2 ml/min. The ACSF was composed of 117 mM NaCl, 4.7 mM KCl, 1.2 mM NaH_2_PO_4_, 2.5 mM CaCl_2_, 1.2 mM MgCl_2_, 25 mM NaHCO_3_, and 11 mM glucose, continuously aerated with a mixture of 95% O_2_ and 5% CO_2_ at pH 7.4.

The neurons were visually patched under a X40 water immersion objective of the microscope (Olympus BX51, Japan) in the whole-cell configuration and were recorded in the current clamp mode. Pyramidal cells in the BLA were identified based on their accommodation properties in response to a sustained depolarizing intracellular current injection. The recording pipettes made from borosilicate glass were filled with a solution containing: 122 mM K-gluconate, 5 mM NaCl, 0.3 mM CaCl_2_, 2 mM MgCl_2_, 1 mM EGTA, 10 mM HEPES, 5 mM Na_2_-ATP, and 0.4 mM Na_2_-GTP, pH adjusted to 7.2–7.3 with KOH (osmolarity adjusted to 280 mOsm/kg with sucrose). The recordings were amplified by a HEKA Patch Clamp EPC10 USB (HEKA Elektronik, USA), analyzed by PatchMaster (HEKA Elektronik, USA) software.

We obtained three dose-response curves to evaluate the effects of U-50488, a kappa opioid receptor agonist, on the spike activity of pyramidal neurons both in the absence and presence of RU-1205. The three experimental conditions included: (1) treatment with U-50488, applied by local perfusion at concentrations ranging from 0.001 to 10 μM (*n*=8); (2) treatment with a combination of U-50488 (0.001–10 μM) and RU-1205 (10 μM) (*n*=8); and (3) treatment with a combination of U-50488 (0.01–10 μM) and RU-1205 (100 μM) (*n*=8). Only one neuron was recorded in each brain slice following the administration of increasing doses of compounds. ACSF served as vehicle control in all experiments.

### Statistical processing

Logistic regression analyses were used to model dose-response curves. Subsequently, key statistical metrics and coefficients were calculated: the maximum effect (E_max_), the coefficient of determination (R^2^) as a statistical measure of model quality, and the slope coefficient which represents the change in the log odds for each unit increase (or decrease) in the predictor variable by 1.0. After verifying the normal distribution with the Shapiro-Wilk test, groups were compared by analysis of variance (ANOVA) with repeated measures followed by Dunnett’s test using GraphPad Prism 10.1. The obtained data are presented as means±SEM. A significance level of p <0.05 was considered significant.

## RESULTS AND DISCUSSION

WPLI values were calculated for 64 electrode pairs and 5 frequency bands (140 parameters for each signal). Taking into account the multicollinearity among the features, further analysis was performed by the principal component analysis in order to reduce the number of variables by combining them into integrative connectivity constructs based on the structure of relationships between them. The first 2 most significant components were identified, which explain 58.32% of the variability (eigenvalues >1).

It has previously been established that the aversive effect of kappa opioid agonists is closely related to the amygdala [17]. Local microinjection of SB203580 (p38 MAPK inhibitor) into the amygdala eliminated the aversive effect of the compound U-50488 (p38 MAPK activator) [18]. Our results showed increased connectivity of the amygdala with various brain regions upon administration of U-50488 (ΔWPLI >0.3 compared to control). Furthermore, these changes in amygdala connectivity were not observed when combining SB203580 with U-50488, thereby confirming an antagonistic interaction between these drugs. Considering that variables associated with the amygdala had high loading coefficients (>0.7) within PC1, it can be assumed that the features related to the effect of drugs on p38 MAPK were captured in this principal component.

The next step was to construct classifier models (Fig. 2) to assess the probability of the compound RU-1205 belonging to the ‘inhibitor’ and ‘non-inhibitor’ classes. To increase stability, predictive reliability, and robustness, a method with a kernel function (Gaussian Process Classifier) was chosen.

**Figure 2.**
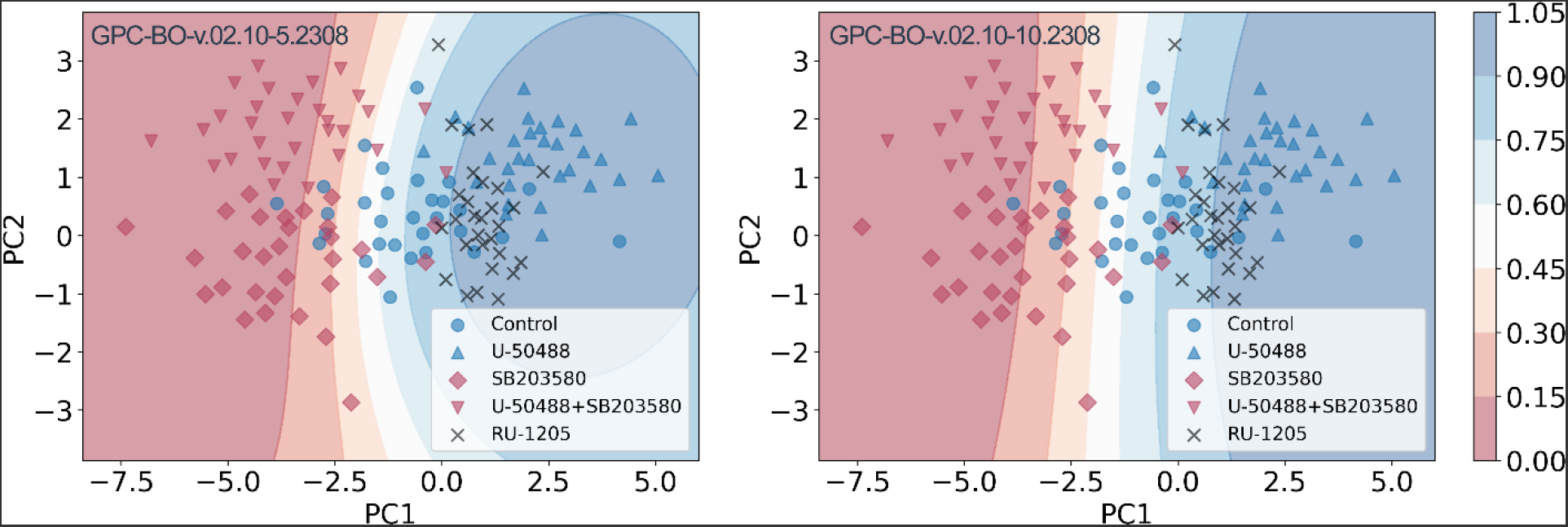
Decision maps of the “Gaussian Process Classifier” (scikit-learn). Note: «Inhibitors» of MAPK p38 are indicated in red, including the subsets «SB203580» and «SB203580+U-50488»; «non-inhibitors» of MAPK p38 are indicated in blue, comprising the ‘control’ subset and the «U-50488» subset. Predicted coordinates for the «RU-1205» signal subset were established. The average probabilities that «RU-1205» signals belong to the «non-inhibitor» class are 0,922584 and 0,894469 as determined by the models «GPC-BO-v.02.10-5.2308» (left) and «GPC-BO-v.02.10-10.2308» (right), respectively.

Compound RU-1205 does not exhibit MAPK p38 inhibitory activity in the central nervous system, as determined with an average probability of at least 89.44% according to the results of the most robust model. To confirm the validity of the models, confusion matrices and the results of 5-fold cross-validation are presented (Fig. 3, 4).

**Figure 3.**
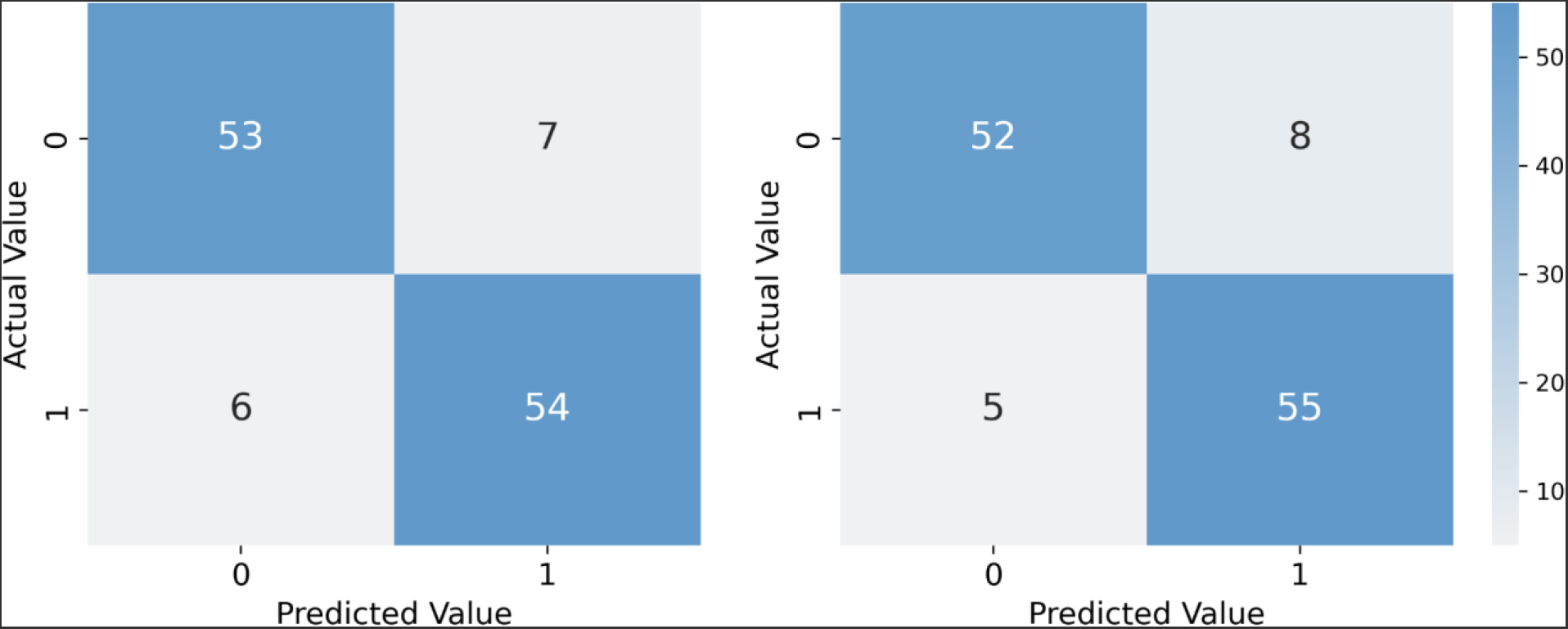
Confusion matrices for the model «GPC-BO-v.02.10-5.2308» (left) and «GPC-BO-v.02.10-10.2308» (right). Note: class 0 – «inhibitors» of p38 MAPK; class 1 – «non-inhibitors» of p38 MAPK.

**Figure 4.**
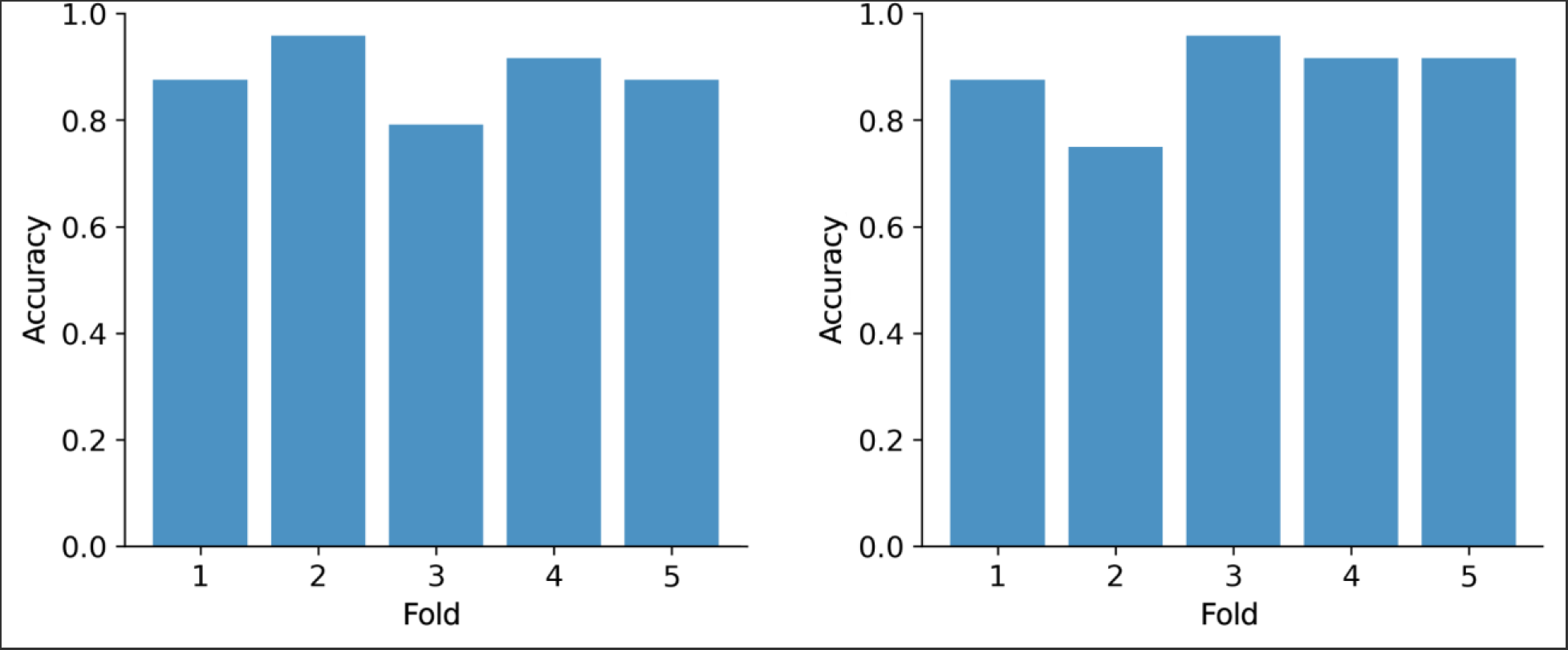
Results of 5-fold cross-validation for two models: «GPC-BO-v.02.10-5.2308» (left) and «GPC-BO-v.02.10-10.2308» (right).

The next part of the work is based on a study which found that blocking kappa opioid receptors in the amygdala leads to a decrease in the spike activity of pyramidal neurons [19]. A similar effect was observed when blocking CRF1 receptors, which, like kappa-opioid receptors, exert their effects through the beta-arrestin pathway and the activation of MAPK p38 [20]. It was also shown that the aversive effect of U-50488 was completely eliminated when the substance SB203580 was injected into the amygdala [18].

To determine the effect of the compound RU-1205 (at a concentration of 10 μM and 100 μM) on U-50488-induced spiking activity of pyramidal neurons in the basolateral amygdala, RU-1205 was applied in combination with increasing concentrations of U-50488.

U-50488 was found to dose-dependently increase the firing rate of BLA pyramidal neurons starting at a concentration of 0.01 μM (p <0.05). The dose-dependent curves (Fig. 5) we observed indicate that the combination of U-50488 and RU-1205 results in competitive antagonism. This is evidenced by a rightward shift in the response curves without a significant change in the slope or maximum effect.

**Figure 5.**
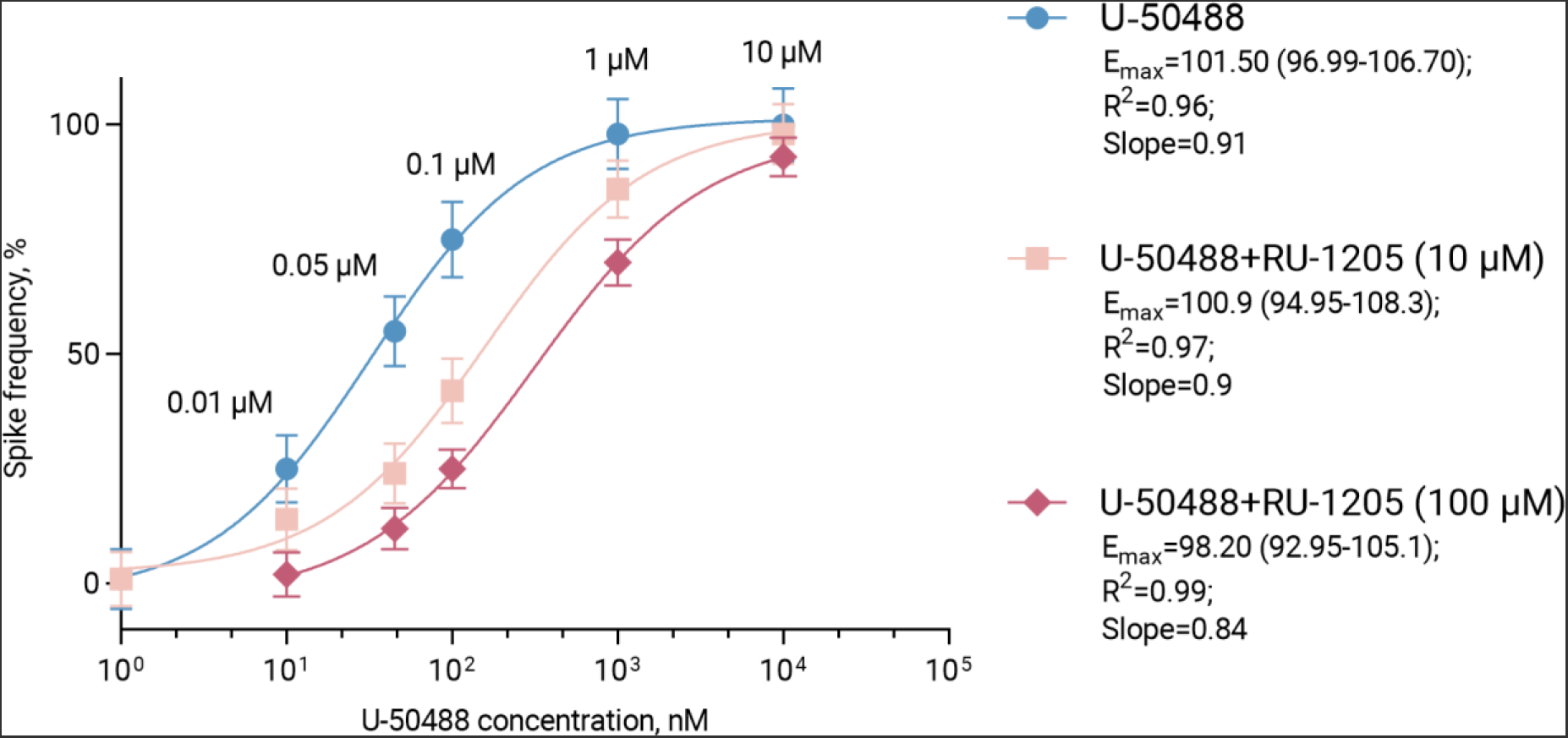
Dose-response relationship of compound U-50488 on the spike frequency of pyramidal neurons in the amygdala, both alone and in combination with different concentrations of compound RU-1205. Note: The graph displays the spike frequency as a function of U-50488 concentration, with curves representing the effect of U-50488 on its own, and in combination with 10 μM and 100 μM of RU-1205. Each data point represents the mean ± SEM.

Various studies have shown that kappa opioid agonists can affect the brain’s bioelectrical activity [21]. The effect of opioid drugs is most pronounced in such brain structures as the cortex [22], hippocampus [23], mesolimbic system [24, 25] and amygdala [26]; therefore, deep electrodes were implanted in these regions. It was previously found that a p38 MAPK inhibitor alters neuronal activity in the hippocampus and also reduces EEG power in the gamma frequency range in mice [27]. Based on this, it was hypothesized that analysis of the effects of kappa opioid agonists and p38 blockers on LFP could provide insight into the mechanism of action of the benzimidazole derivative RU-1205 with a similar activity profile. Given the nonspecific nature of spectral changes in LFP, we attempted to extract features of pharmacological action from a set of WPLI functional connectivity matrices. WPLI was chosen as a method for quantifying connectivity due to its higher resistance to volume conduction and lower sensitivity to noise [28]. WPLI is widely used in electroencephalographic (EEG) research [29], electrocorticographic (ECoG) studies [30], and LFP analysis, including studying the effects of substances with psychotropic activity [31].

To further analyze the resulting WPLI scores after administration of p38 MAPK inhibitors and kappa opioid agonists, a combined approach was adopted, including principal component analysis and construction of a Gaussian process regression model for signal classification. Principal component analysis is used in signal processing and machine learning to reduce the dimensionality of data while preserving as much information as possible. The performance of the method, when combined with various classifier models, has been confirmed in EEG studies [32]. The Gaussian process regression model has also shown high efficiency (94% accuracy achieved for three classes) in analyzing the features of EEG signals associated with stress levels [33], which seems important since kappa opioid agonists are distinguished by their ability to provoke stress and anxiety.

The advantage of the proposed approach is that nervous tissue is an ideal detector of a neuro-or psychotropic substance, since its effects are specifically reflected in bioelectrical activity, the decoding of which makes it possible to construct highly accurate predictions. Modern analytical methods make it possible to identify more subtle and complex signal patterns associated with pharmacological action. This is evidenced by the consistent growth in scientific interest and the rising number of studies focusing on automated EEG classification [34, 35]. One significant area of focus involves classifying EEG, ECoG, and LFP signals to discern the psychotropic effects and elucidate the mechanisms of action of test substances [36, 37].

To verify the reliability of the findings regarding the mechanism of action of the compound RU-1205, additional studies were carried out using the patch clamp method on acute slices of the rat brain. The presented results allow us to conclude that the compound RU-1205 does not exhibit activity similar to SB203580, but competes for a common binding site with the compound U-50488, thus eliminating its aversive effect, an indirect sign (correlate) of which, within this model, is increased spike activity of neurons in the basolateral complex of the amygdala. This finding is a hallmark of functional selectivity. Specifically, the functional agonist 6′-GNTI has been shown to reduce the arrestin-mediated effects of unbiased agonists [38]. At the same time, when combining the compounds U-50488 and SB203580, non-competitive antagonism was observed, as evidenced by a change in the slope coefficient and a reduction in the maximum effect of U-50488 (p <0.05) [39].

### Limitations of the study

It should be noted that the approach used to classify LFP has a number of limitations. A primary concern is volume conductivity. Electrical signals generated by neurons can propagate in brain tissue, leading to distortion and mixing of signals from different sources. It should also be taken into account that in many brain structures most of the LFP activity originates from remote current sources rather than ones local to the electrode [40].

In addition, the predictive accuracy of machine learning models drops significantly when extrapolating beyond the training data, a scenario known as **«**out-of-distribution». Therefore, our training dataset included additional signals obtained after the administration of the combination **«**SB203580+U-50488», which simulates a dual mechanism of action. However, this did not completely eliminate the identified problem.

Conducting a study with repeated measures, where different drugs are tested on the same group of animals, has several disadvantages, including carryover effects, in which the effect of one drug persists and influences subsequent responses, and order effects, where the sequence of drug administration significantly affects the results. Psychological conditioning can be associated with changes in animals’ responses over time, potentially skewing the results. This study design may have limited generalizability due to the homogeneous pool of subjects when extrapolating results to the general population, but, on the other hand, it eliminates the contribution of interclass variability and allows for better performance of the classifier. In addition, when substances are administered sequentially, the need to wait for their complete (or near-complete) elimination not only prolongs the study’s duration but also heightens the risk of the animal being excluded from the experiment due to the development of complications or death.

## CONCLUSION

The study presents a comprehensive approach to analyze brain connectivity, which includes recording electrophysiological data and using machine learning methods to classify pharmacological compounds based on LFP changes. We identified integrative connectivity characteristics derived from WPLI calculations and principal component analysis. These characteristics were associated with the effects of kappa opioid agonists and p38 MAPK inhibitors. The Gaussian process classifier allowed us to classify the compound RU-1205 as a «non-inhibitor» of p38 MAPK with a probability of 0.89.

The results obtained were confirmed in patch clamp experiments on living brain slices. It was demonstrated that RU-1205 interacts with U-50488, competitively suppressing its effect on the spike activity of BLA pyramidal neurons. In contrast, the p38 MAPK inhibitor SB203580 non-competitively inhibits spike activity induced by U-50488. This suggests that compound RU-1205 exhibits functional kappa agonist activity and does not have a significant effect on p38 MAPK.

The study illustrates the potential for combining electrophysiological measurements with advanced data analysis methods to gain a comprehensive understanding of the neuronal mechanisms of action, and also highlights the promise of further research in this direction.

## Abbreviations

BBB: blood-brain barrier;
CNS: central nervous system;
ACSF: artificial cerebrospinal fluid;
BLA: basolateral amygdala;
LFP: local field potential;
p38 MAPK: p38 mitogen-activated protein kinase;
WPLI: weighted phase lag index.

## ACKNOWLEDGMENTS

The authors express their gratitude to the team of the Institute of Physical and Organic Chemistry at Southern Federal University for the synthesis of the studied compound RU-1205.

## AUTHORS’ CONTRIBUTION

All authors confirm that their authorship meets the international ICMJE criteria (all authors made a significant contribution to the development of the concept, conduct of the study and preparation of the article, read and approved of the final version before the publication). K.Y. Kalitin -statement of key objectives, analysis of scientific and methodical literature, data processing, writing, and editing of the manuscript; O.Y. Mukha - data collection, data processing, writing, editing, and formatting of the manuscript; A.A. Spasov - critical revision of draft manuscript with valuable intellectual investment, final manuscript approval.

## Footnotes

1 Zenodo. MNE-Python (v1.6.1). - [Electronic resource]. – Access mode: https://zenodo.org/records/105199482

2 Scikit-learn. Machine Learning in Python. - [Electronic resource]. – Access mode: https://scikit-learn.org/stable/index.html

